# HistoSB-Net: Semantic Bridging for Data-Limited Cross-Modal Histopathological Diagnosis

**DOI:** 10.64898/2026.03.23.713838

**Authors:** Bingyuan Bai, Tzu-Ching Shih, Kazunori Miyata

## Abstract

Vision–language models (VLMs) provide a unified framework for multimodal reasoning, yet their representations are primarily learned from natural image–text corpora and often exhibit semantic misalignment when transferred to histopathology, particularly under data-limited diagnostic settings. To address this limitation, we propose HistoSB-Net, a semantic bridging network designed to adapt pre-trained VLMs to multimodal histopathological diagnosis while preserving their original semantic structure. HistoSB-Net introduces a constrained semantic bridging (CSB) module that operates within the self-attention projection space of both vision and text encoders. Instead of employing explicit cross-attention or full fine-tuning, CSB adaptively modulates pre-trained attention projections through a lightweight nonlinear semantic bottleneck, enabling structured cross-modal regulation with limited additional parameters. The framework supports both patch-level and whole-slide image (WSI)-level diagnosis within a unified architecture. Experiments on six pathology benchmarks, comprising two WSI-level and four patch-level datasets, demonstrate consistent improvements over zero-shot inference across 36 backbone–dataset combinations under limited supervision. Further analysis of prototype-based margin distributions and confusion matrices shows that these improvements are accompanied by enhanced intra-class compactness and increased inter-class separation in the embedding space. These results indicate that CSB provides an effective and computationally manageable strategy for adapting pre-trained VLMs to data-limited digital pathology tasks.

## 1. Introduction

Histopathological examination remains the gold standard for definitive cancer diagnosis and prognostication [7]. With the digitization of whole-slide images (WSIs) [9], computational pathology (CPath) has enabled large-scale analysis of high-resolution tissue morphology, revealing phenotypic characteristics of the tumor microenvironment that reflect tumor aggressiveness and biological behavior [2]. However, compared with natural image benchmarks, pathology datasets typically require expert-level clinical annotations and ethical approval, resulting in limited availability of labeled data [19]. In few-shot scenarios, this scarcity constrains model adaptation to substantial intra-class heterogeneity and subtle inter-class differences, thereby complicating robust representation learning [24].

In contrast to traditional unimodal recognition paradigms such as convolutional neural networks, vision–language models (VLMs), exemplified by Contrastive Language– Image Pre-training (CLIP) [26], align visual and textual encoders in a shared embedding space through large-scale contrastive pre-training. By leveraging natural language supervision, VLMs enable zero-shot inference via prompt-based matching, reducing reliance on labeled data and offering an alternative to task-specific classifier training.

However, directly transferring VLMs to pathology introduces substantial challenges. A prominent example arises from the pervasive intra-class heterogeneity and inter-class homogeneity in histopathological images (Fig. 1(a)–(b)) [32], where substantial visual variability exists within the same diagnostic category while distinct categories may exhibit overlapping tissue patterns. Under such conditions, generic textual prompts often fail to provide sufficiently discriminative semantic guidance, as subtle morphological differences cannot be reliably captured by fixed template-based descriptions. A straightforward approach to mitigate this domain gap is prompt refinement through contextual or descriptive templates. While prompt refinement can improve zero-shot performance, it typically relies on heuristic trial-and-error design and adjusts only the textual input without modifying the internal feature representations of the model, resulting in limited adaptability [20].

**Figure 1.**
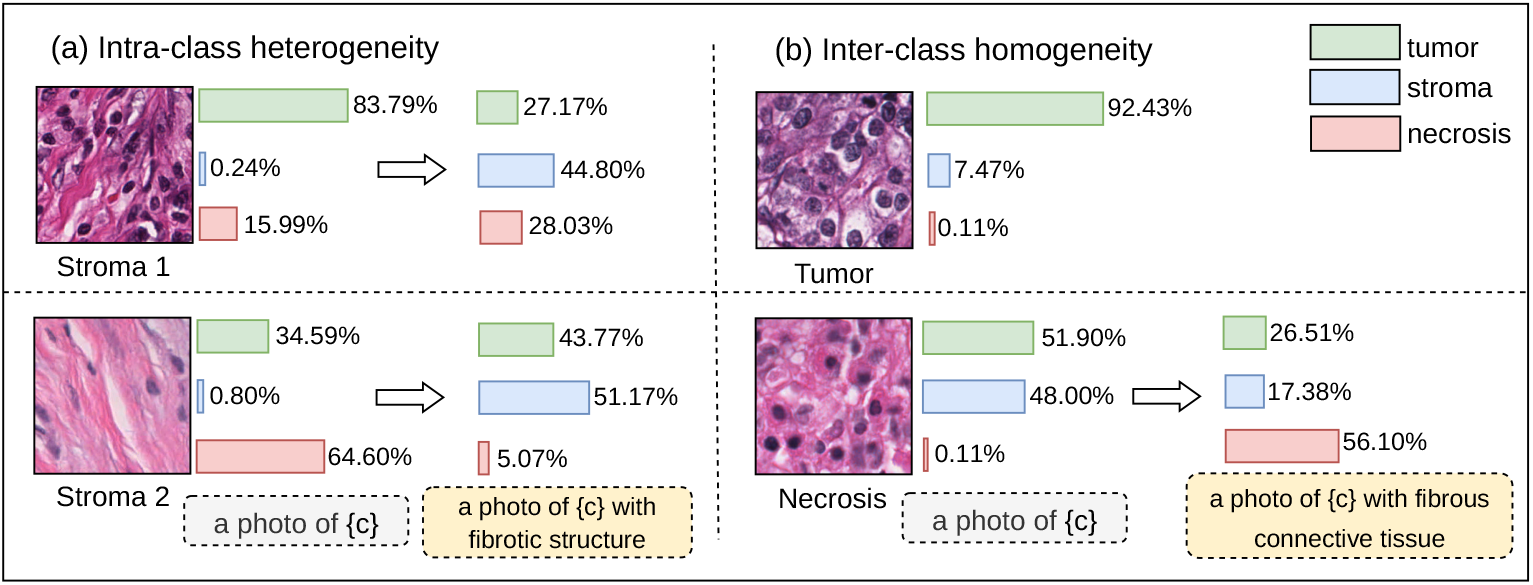
Effect of prompt refinement on zero-shot classification in histopathology. (a) Under pronounced intra-class heterogeneity, two image patches belonging to the stroma class are incorrectly predicted when using a generic prompt template. (b) Under inter-class homogeneity, a patch from the necrosis class is misclassified as tumor with the same generic prompt. In both cases, incorporating class-relevant contextual information into the prompts corrects the predictions. All zero-shot classification results are obtained using the OpenAI pre-trained CLIP ViT-B/16 backbone.

**Figure 2.**
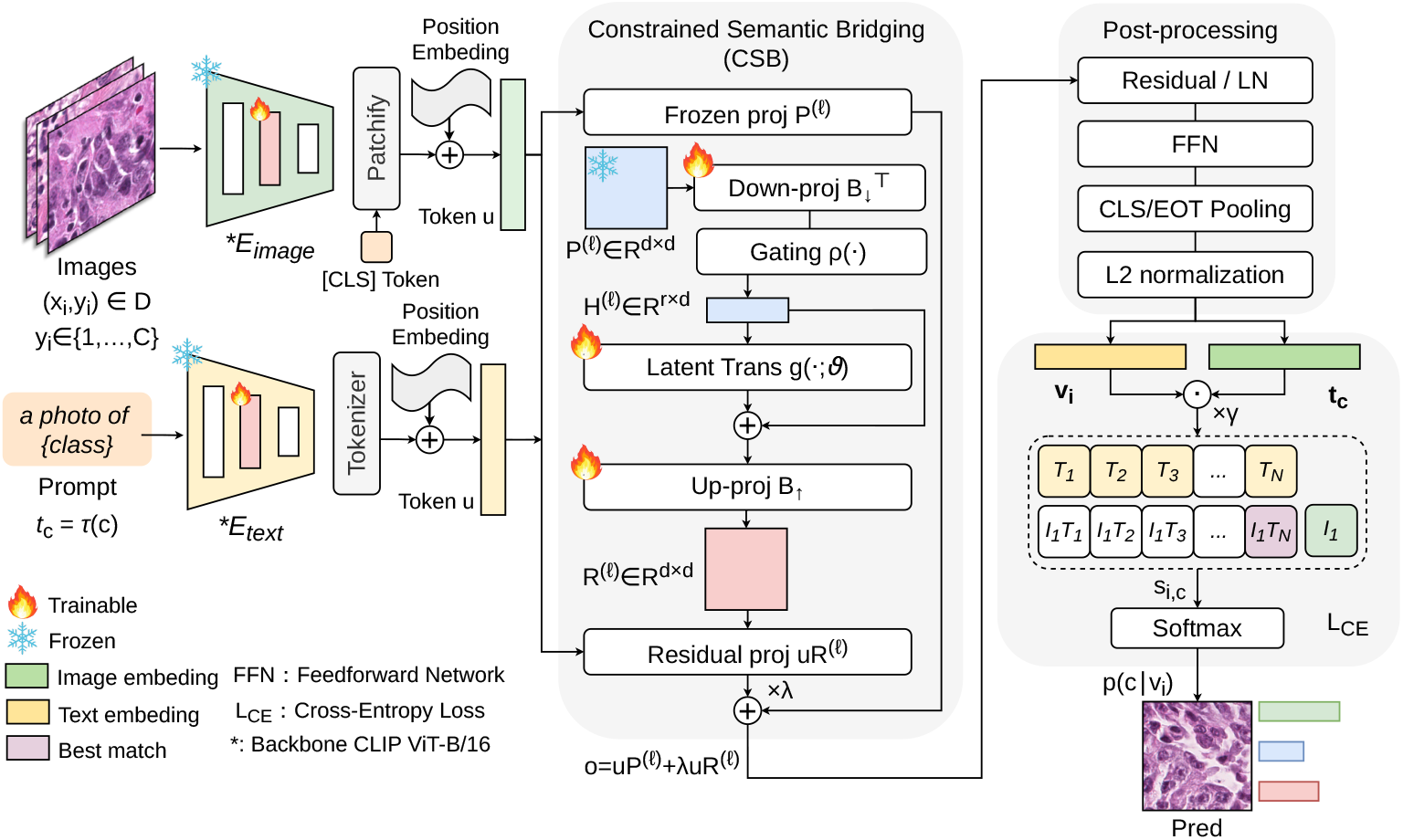
Overview of HistoSB-Net. Given an image *x*_*i*_ and a class prototype text *t*_*c*_ = *τ*(*c*), the frozen CLIP ViT-B/16 vision and text backbones produce embeddings with CSB enabled on selected self-attention projections. CSB generates a projection residual from each frozen attention projection and injects it as a scaled additive modulation while keeping all backbone weights fixed. The resulting *ℓ*_2_-normalized embeddings are matched via temperature-scaled cosine similarity and optimized using cross-entropy over the ground-truth labels.

To enhance the few-shot adaptation capability of pre-trained VLMs under limited supervision, various lightweight adaptation strategies have been proposed, including prompt learning and adapter-based tuning. These approaches offer clear computational advantages by avoiding full fine-tuning, yet they still encounter challenges in CPath, where structural complexity and subtle inter-class differences are prominent.

Representative prompt learning such as CoOp [43] replace manually designed templates with learnable continuous context vectors while keeping the backbone frozen. However, their optimization is confined to input-level reparameterization on the textual side, and the learned continuous prompts exhibit limited interpretability and sensitivity to label noise. Adapter-based methods, exemplified by CLIP-Adapter [10], introduce lightweight bottleneck modules with residual blending to adjust feature representations. Although effective in few-shot settings, their performance remains sensitive to design choices such as the residual ratio, requiring a balance between adaptation strength and stability.

Recently, low-rank adaptation (LoRA) [14] models fine-tuning as additive low-rank perturbations to frozen pre-trained weights and has demonstrated that attention projection layers constitute effective loci for efficient adaptation in transformer-based models. In vision–language settings, attention projections determine how token embeddings are linearly transformed prior to similarity computation, thereby structuring the space in which cross-modal alignment emerges. Prior work has reported performance degradation and cross-domain generalization challenges when transferring VLMs from natural image–text corpora to medical pathology data under domain shift [19]. Given that projection layers govern how embeddings are organized before similarity estimation, regulating projection transformation becomes particularly relevant under severe distribution shifts.

This observation motivates a complementary perspective: beyond modifying model parameters, can adaptation be achieved by directly regulating how projection layers transform token embeddings, thereby reshaping the interaction geometry within which cross-modal alignment unfolds?

To investigate this perspective, we propose HistoSB-Net, a semantic bridging framework for CPath cross-modal learning that performs projection-aware adaptation at the computation level. Instead of altering backbone weights, we introduce structured residual modulation within the projection computation itself. Specifically, we design a Cross-Modal Semantic Bridging (CSB) module that derives a compact latent transform from frozen attention projections and injects it as a controlled additive residual into selected projection layers of both the vision and text encoders. By modulating the projection-induced embedding transformation while preserving all backbone parameters, CSB reshapes the token embedding geometry underlying self-attention, influencing how interactions emerge without modifying the attention mechanism. For model optimization, we follow the standard supervised classification objective of the downstream task: given the CSB-modulated visual and textual embeddings, classification is performed by temperature-scaled cosine similarity, and the model is trained using standard cross-entropy on cosine-similarity-based logits.

The contributions are summarized as follows:

- **A projection-aware adaptation framework**. We propose HistoSB-Net, a projection-level adaptation framework that performs structured modulation within attention projections for multimodal histopathology diagnosis. Under 16-shot supervision, HistoSB-Net achieves Macro-F1 scores of 82.34% on BCSS, 83.66% on GCSS, 85.70% on BCSS-WSSS, 81.60% on LUAD-HistoSeg, 72.11% on EBHI-Seg, and 84.17% on PathM-NIST, consistently outperforming zero-shot inference and representative lightweight adaptation baselines across six benchmarks.
- **CSB module**. We introduce a CSB module that injects a structured residual transformation into selected attention projections of both vision and text encoders while keeping the backbone fully frozen. The module accounts for only 0.49% of the total ViT-B/16 parameters and maintains stable computational cost, with average training time per epoch between 37.40 s and 48.00 s and GPU memory usage below 22.39% of a 24 GB RTX 4090.
- **Representation-level separability improvement**. Beyond performance gains, we demonstrate that HistoSB-Net improves embedding geometry. Across all datasets, the average class discriminability margin increases substantially, for example from 0.010 to 0.083 on BCSS and from 0.014 to 0.130 on LUAD-HistoSeg, accompanied by consistent increases in the proportion of positively separated samples. Confusion matrix analysis further shows strengthened diagonal dominance and reduced inter-class overlap, indicating improved intra-class compactness and inter-class separation under limited supervision.

## 2. Related work

This section reviews recent vision–language modeling approaches for CPath, with an emphasis on their cross-modal alignment mechanisms, supervision sources, and data regimes. We further highlight the limitations of these methods in data-limited diagnostic settings.

### 2.1. VLMs for CPath

Recent research on VLMs for CPath has largely focused on large-scale cross-modal alignment through data construction and contrastive pretraining. A major research line builds extensive paired pathology image–text datasets to support CLIP-style contrastive learning. For example, PathText provides nearly 10,000 WSI–text pairs and has been used to train generative models such as MI-Gen [4], while PathGen-1.6M [31] offers over one million image–caption pairs for large-scale pretraining and evaluation. Quilt-1M [16] further curates large-scale pathology image–text pairs via automatic speech recognition (ASR), enabling CLIP-style fine-tuning at scale. In this paradigm, models such as PLIP [15] and PathCLIP adapt CLIP to pathology by optimizing global embedding alignment between visual patches and textual descriptions, while CHIEF [33] strengthens visual representations prior to alignment through Transformer-based encoders. KEEP [44] extends this strategy by incorporating structured pathology knowledge graphs to guide contrastive pair construction beyond simple image–text matching.

Beyond pure contrastive alignment, another research direction integrates generative supervision following the contrastive captioning (CoCa) formulation [38]. CONCH combines self-supervised visual pretraining with CoCa-style joint optimization to support both retrieval and caption generation in pathology scenarios, while PRISM [27] extends multimodal alignment to WSI and report-level modeling through long-range context encoding and structured report generation. TITAN [8] further integrates self-supervised visual encoders with CoCa training for multi-scale pathology representation learning, and MUSK [35] explores hybrid objectives that fuse large-scale masked modeling on unpaired data with subsequent CoCa-style multimodal contrastive training on paired data to enhance robustness and generalization. These generative paradigms enrich multimodal supervision but typically confine cross-modal interaction to shared embedding alignment or late-stage fusion.

In parallel, several studies incorporate VLMs into down-stream WSI modeling pipelines. FSWC [25] leverages CLIP features and employs GPT-4 to generate grouped prompts corresponding to pathological sub-concepts for weakly supervised WSI classification, while ViLA-MIL [29] introduces a language-augmented multiple-instance learning framework in which GPT-3.5 generates multi-scale textual descriptions fused with visual representations. HLSS [34] proposes hierarchical semantic supervision to align visual features with multi-level diagnostic labels and textual semantics, and CPLIP [17] constructs a pathology-specific vision–language dictionary with many-to-many contrastive learning and prompt transformations derived from MI-Zero [22] and GPT-based rewriting. Visual question answering benchmarks such as PathVQA [13] and WSI-VQA [5] further extend multimodal supervision toward fine-grained diagnostic reasoning scenarios.

Despite these advances, most existing VLM-based methods for CPath primarily rely on data scale or external language priors to achieve alignment. In pathology scenarios characterized by limited data, sparse supervision, and significant domain shifts, such strategies may lead to unstable alignment performance[19]. This limitation motivates the development of adaptation mechanisms to enable efficient domain transfer.

### 2.2. Parameter-efficient adaptation

Existing adaptation strategies differ primarily in where structural intervention is introduced within the model. Prompt-based methods extend continuous context learning in various directions. CoCoOp [42] introduces instance-conditional prompts to improve generalization, while ProDA [23] models prompts as distributions to capture diversity in few-shot regimes. KgCoOp [37] incorporates knowledge-guided regularization to prevent drift from pre-trained priors, ProGrad [45] constrains gradient updates to preserve general knowledge, and PLOT [3] stabilizes multi-prompt optimization via optimal transport-based alignment. These approaches operate primarily through input-level reparameterization on the textual or visual side.

Another line of work introduces lightweight modules or auxiliary structures to adjust intermediate representations in feature space. Tip-Adapter [41] constructs a key–value cache from few-shot features for training-free or minimally fine-tuned adaptation, while TaskRes [39] adds residual task-specific parameters to classifier weights to decouple down-stream knowledge from pre-trained priors. Vision prompt tuning [18] injects learnable visual tokens into transformer inputs without modifying backbone weights, and AdaptFormer [6] integrates bottleneck-style adapters within transformer blocks for efficient adaptation of vision backbones. These methods primarily operate in feature or classifier space by introducing auxiliary modules while preserving the original architecture.

As summarized in Table 1, existing lightweight adaptation strategies primarily operate by refining textual prompts or inserting auxiliary feature-level modules. While effective under moderate distribution shifts, these approaches either confine adaptation to the input side or adjust representations only after encoder computation, providing limited control over how projection layers shape cross-modal embedding transformation, which becomes particularly critical under severe domain shifts such as those encountered in histopathology. To address this limitation, recent work has begun to explore adaptation directly at the projection layers. Building on the LoRA mechanism, CLIP-LoRA [40] applies low-rank updates to attention projection layers in VLMs, demonstrating that projection layers serve as effective loci for few-shot adaptation with limited additional parameters.

**Table 1.** Comparison of representative VLM adaptation strategies.

| Example | Key Tech | Intervention | Advantages | Limitations |
| --- | --- | --- | --- | --- |
| CoOp | Learnable context tokens | Text input | Avoids manual prompt design | Limited control over visual branch |
| CLIP-Adapter | Residual feature adapters | Feature space | Direct feature-level adaptation | Sensitive to training settings |
| HistoSB-Net (Ours) | Projection residual modules | Attention projection | Regulates representation formation | Introduces additional computation |

Inspired by this observation, HistoSB-Net explores an alternative perspective on projection regulation. Instead of modifying projection weights through low-rank perturbations, the proposed framework introduces structured residual modulation within the projection computation itself. A residual transform is derived from frozen projections and injected at the projection output, reshaping the projection-induced embedding transformation while preserving the original attention mechanism. In this way, adaptation shifts from parameter perturbation around a fixed mapping to computation-level geometric regulation.

## 3. Methods

### 3.1. Data organization

Given a pathology dataset 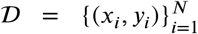, each sample consists of an image *x*_*i*_ and a categorical label *y*_i_ ∈ {1, …, *C*}, where *C* denotes the number of diagnostic categories. We consider a few-shot learning setting in which each category *c* is associated with *K* labeled training samples. For each category *c*, a textual class description *t*_*c*_ is constructed using a fixed prompt template,

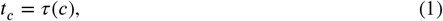

where *τ*(⋅) denotes a fixed generic text template of the form “a photo of {}”, and the resulting class description is shared across all samples belonging to category *c*.

### 3.2. CSB

As shown in Fig. 3, CSB regulates adaptation by conditioning a frozen attention projection on its own geometry through a compact semantic transform, and then injecting the resulting modulation as a residual update at the projection output.

**Figure 3.**
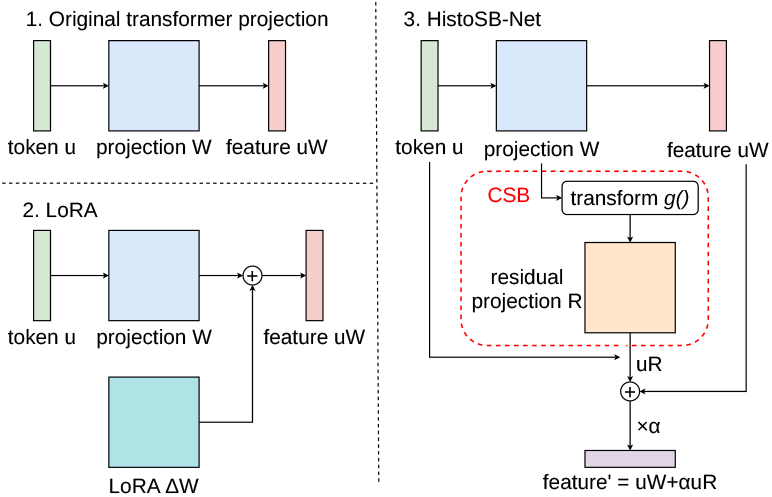
Comparison of the original projection, LoRA weight update, and CSB residual modulation.

Let **P**^(*ℓ*)^ ∈ ℝ^*d*×*d*^ denote a frozen linear projection in the *ℓ*-th self-attention block. CSB first extracts a compressed projection representation from **P**^(*ℓ*)^ via a learned contraction,

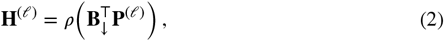

where **B**_↓_ ∈ ℝ^*d*×*r*^ performs a contraction along the input dimension, producing **H**^(*ℓ*)^ ∈ ℝ^*r*×*d*^. Here *r* controls the contraction dimension used to summarize the frozen projection, rather than enforcing a low-rank constraint on **P**^(*ℓ*)^ itself. *ρ*(⋅) denotes the Gaussian Error Linear Unit (GELU) activation applied element-wise.

A lightweight residual refinement is then applied in this compressed space to capture structured semantic variation while preserving the coarse geometry induced by the pretrained projection. Concretely, *g*(⋅; *ϑ*) is a shallow latent transform parameterized by *ϑ*,

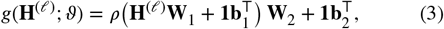

where *ϑ* = {**W**_1_, **b**_1_, **W**_2_, **b**_2_}, **W**_1_ ∈ ℝ^*d*×*m*^, **W**_2_ ∈ ℝ^*m*×*d*^, **b**_1_ ∈ ℝ^*m*^, **b**_2_ ∈ ℝ^*d*^, and **1** is an all-ones column vector for bias broadcasting. The refined compressed representation is computed as

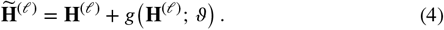

The refined code is lifted back to the original projection space to form a projection residual,

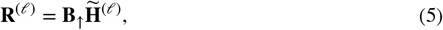

with **B**_↑_ ∈ ℝ^*d*×*r*^. Since **R**^(*ℓ*)^ is explicitly derived from the frozen projection **P**^(*ℓ*)^, the induced modulation remains projection-conditioned and preserves the structural prior encoded in the pre-trained weights.

Given an input token representation **u**, CSB modulates the projection output by injecting this residual in a scaled additive form,

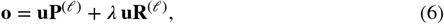

where *λ* is a fixed hyperparameter controlling the strength of projection-level modulation. The same CSB parameterization is shared across all samples and classes, and is applied symmetrically to both the vision and text branches.

In implementation, CSB is activated only on designated self-attention blocks. Let 𝒮_*v*_ and 𝒮_*t*_ denote the selected layer index sets for the vision and text en^*t*^ coders, respectively, and let 𝒜 = {*q, k, v, o*} denote the attention projections. For each branch *b* ∈ {*v, t*}, CSB is applied as

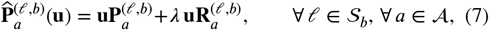

and reduces to the frozen projection otherwise,

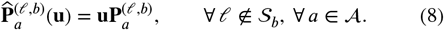

Equivalently, using an indicator 𝕀[⋅],

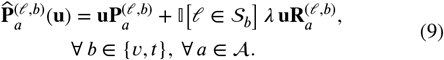

According to the layer selection described above, CSB is instantiated by replacing the attention projections in the designated blocks with 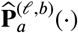 for *ℓ* ∈ 𝒮_*b*_. Each branch then produces a sequence of CSB-modulated^*b*^token features, from which a global embedding is obtained via the standard backbone readout. Specifically, for the vision encoder, the representation corresponding to the [CLS] token is used as the global visual embedding, while for the text encoder, the embedding at the end-of-text token position is used as the global textual representation. The final visual and textual representations used for training are

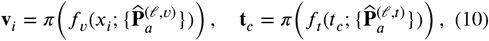

where the original backbone parameters remain frozen and only CSB parameters are updated.

### 3.3. Training objective

Under the CSB formulation described above, model adaptation is achieved by modulating a subset of frozen attention projections through the induced operators 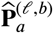, while all original backbone parameters remain fixed. Let *f*_*v*_(⋅; *ϑ*) and *f*_*t*_(⋅; *ϑ*) denote the vision and text encoders with CSB-modulated projections applied at the selected layers, where *ϑ* collects all learnable CSB parameters. For an input image *x*_*i*_ and class text prototype *t*_*c*_, the corresponding global representations are obtained through the standard backbone readout operator *π*(⋅) as

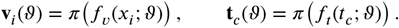

Both embeddings are *ℓ*_2_-normalized. Classification is performed by computing the cosine similarity

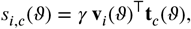

where *γ* is a fixed scaling factor. The similarity vector **s**_*i*_(*ϑ*) serves as the logit vector for the downstream classification task. The CSB parameters *ϑ* are optimized by minimizing the standard supervised cross-entropy loss over the training distribution 𝒟,

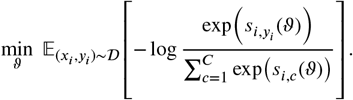

For evaluation, we use macro-averaged F1 to reflect balanced recognition across diagnostic prototypes. Predictions are obtained from the cosine-similarity logits,

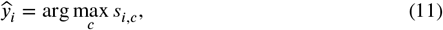

where *s*_*i,c*_ is the temperature-scaled normalized similarity between **v**_*i*_ and the prototype **t**_*c*_. Based on 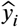 and the ground-truth label *y*_*i*_, we compute TP_*c*_, FP_*c*_, and FN_*c*_ for each prototype *c*. The evaluation metric is then

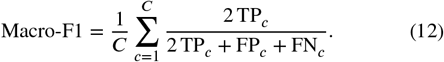

## 4. Datasets and training setting

We conducted experiments on five publicly available hematoxylin and eosin (H&E)–stained histopathology benchmarks, encompassing both WSI and patch-level classification tasks.

Following the prototype construction strategy of Tang et al. [32], we build a pathology-specific vision–text semantically aligned prototype library for each benchmark as the foundation for subsequent data-limited training (Table 2). Each prototype represents a category-aware visual instance paired with its textual label, designed to preserve representative semantic patterns within each class.

**Table 2.** Statistics of the prototype images for each benchmark, including dataset type, tissue category, total number of prototypes, and number of classes.

| Dataset | Type | Tissue | Number | Classes |
| --- | --- | --- | --- | --- |
| BCSS[1] | WSI | Breast | 13,932 | 4 |
| GCSS[28] | WSI | Stomach | 274 | 5 |
| BCSS-WSSS[11] | Patch | Breast | 9,378 | 4 |
| LUAD-HistoSeg[12] | Patch | Lung | 4,595 | 4 |
| EBHI-Seg[30] | Patch | Colon | 2,228 | 6 |
| PathMNIST[36] | Patch | Colon | 1,600 | 8 |

For the two WSI-level datasets, a two-stage preprocessing strategy is adopted. We first follow the protocols described in the original studies to exclude non–region-of-interest areas while retaining the slide-level organization. The resulting whole-slide images are then partitioned into tissue patches following standard VLM-based WSI classification pipelines [21]. For the patch-level benchmarks, images are directly grouped by category. After this pre-processing, all samples are unified at the patch level and organized into class-specific prototype pools.

Under the data-limited setting, 16 images per class are randomly sampled from the prototype pool using a fixed seed to ensure reproducibility. These samples are split into training, validation, and test sets with a ratio of 8:1:1. During training, the checkpoint achieving the best validation Macro-F1 is selected, and its performance is subsequently evaluated on the full test set of each benchmark. All experiments are conducted under a unified and reproducible training configuration to ensure fair comparison across methods, with the key hyperparameters summarized in Table 3.

**Table 3.** Training configuration and hyperparameter settings used in all HistoSB-Net experiments.

| Component | Configuration |
| --- | --- |
| Operating system | Ubuntu 24.04 LTS |
| GPU | 1 NVIDIA RTX 4090 |
| Deep learning framework | PyTorch |
| Backbone | CLIP ViT-B/16 |
| Text prompt | "a photo of {c}" |
| Learning rate (LR) | $2 \times 10^{-3}$ |
| LR schedule | Cosine decay |
| Optimizer | AdamW |
| Batch size | 8 |
| Gradient accumulation | 4 steps |
| Gradient clipping | Max-norm 1.0 |
| Dropout rate | 0.1 |
| Samples per class (shots) | 16 |
| Contraction dimension $r$ | 8 |
| Scaling factor $\lambda$ | 2 |
| Regulated encoder branches | Vision and text |
| Projection type | q, k, and v |
| Random seed | 1, 2, 3, 4, 5 |
| Number of training epochs | 50 |

## 5. Experiments

This section presents a systematic empirical evaluation of HistoSB-Net under the setting described in Table 3. The experimental analysis is organized into five parts corresponding to the subsequent subsections. (1) **Zero-shot comparison**. Quantitative evaluation against zero-shot inference across diverse pre-trained vision–language backbones examines whether CSB consistently enhances cross-modal diagnostic alignment under data-limited supervision. (2) **Adaptation comparison**. Performance is bench-marked against representative prompt-based, adapter-based and CLIP-LoRA methods to assess effectiveness across adaptation paradigms. (3) **Ablation analysis**. Controlled experiments analyze the influence of supervision scale, encoder branch selection, and projection insertion strategies on adaptation behavior. (4) **Class discriminability**. Prototype-based margin distributions and confusion matrices are used to investigate whether performance gains correspond to improved intra-class compactness and inter-class separation in the embedding space. (5) **Computational cost**. Training overhead and parameter efficiency are reported to quantify the trade-off between diagnostic performance and computational complexity.

### 5.1. Zero-shot baseline

This subsection evaluates whether CSB improves diagnostic performance beyond frozen pre-trained representations and whether such improvements generalize across diverse VLM backbones. We first compare HistoSB-Net with standard zero-shot inference under the prompt template “a photo of {c}”, using multiple pre-trained VLMs including CLIP ViT-B/16, CLIP ViT-L/14, BiomedCLIP, PLIP, MI-Zero, and Quilt-1M.

We then integrate the proposed CSB module into these pre-trained backbones to assess whether it consistently enhances performance across different encoder architectures and pretraining corpora. All results are reported in Table 4.

**Table 4.** Macro-F1 (%) comparison between zero-shot baselines of different pre-trained VLMs and their performance after CSB-based adaptation.

| Model | BCSS | GCSS | BCSS-WSSS | LUAD-HistoSeg | EBHI-Seg | PathMNIST |
| --- | --- | --- | --- | --- | --- | --- |
| *Zero-shot Inference* |  |  |  |  |  |  |
| CLIP ViT-B/16 (Baseline) | 11.41 | 16.34 | 15.64 | 16.46 | 16.93 | 10.10 |
| CLIP ViT-L/14 (ICML 2021) | 12.83 | 35.49 | 14.39 | 33.11 | 22.37 | 28.99 |
| BiomedCLIP (NeurIPS 2023) | 21.34 | 36.55 | 23.15 | 50.86 | 18.41 | 24.71 |
| PLIP (Nature Medicine 2023) | 27.21 | 36.39 | 28.16 | 54.67 | 29.23 | 53.99 |
| MI-Zero (CVPR 2023) | 18.16 | 24.13 | 18.73 | 21.28 | 5.69 | 7.66 |
| QuiltNet (NeurIPS 2025) | 20.96 | 40.40 | 24.15 | 42.83 | 19.13 | 25.20 |
| *CSB Module* |  |  |  |  |  |  |
| CLIP ViT-B/16 (HistoSB-Net) | 82.34 ± 3.59 | 83.66 ± 3.72 | 85.70 ± 3.71 | 81.60 ± 2.69 | 72.11 ± 3.15 | 84.17 ± 5.59 |
| CLIP ViT-L/14 | 83.31 ± 3.25 | 87.35 ± 3.36 | 86.48 ± 2.81 | 83.20 ± 5.73 | 73.32 ± 2.83 | 87.46 ± 2.63 |
| BiomedCLIP | 77.12 ± 1.28 | 83.40 ± 1.91 | 85.51 ± 2.77 | 84.51 ± 3.85 | 69.00 ± 5.60 | 85.76 ± 3.13 |
| PLIP | 81.11 ± 4.32 | 82.61 ± 10.26 | 82.89 ± 8.24 | 86.35 ± 2.75 | 72.19 ± 4.09 | 86.50 ± 5.57 |
| MI-Zero | 74.23 ± 5.25 | 83.89 ± 5.25 | 73.49 ± 3.46 | 74.38 ± 7.20 | 51.29 ± 7.17 | 72.35 ± 9.67 |
| QuiltNet | 75.07 ± 8.47 | 80.90 ± 4.20 | 83.67 ± 3.94 | 84.74 ± 11.35 | 72.74 ± 1.97 | 85.38 ± 5.92 |

As shown in Table 4, integrating the CSB module yields substantial and consistent performance improvements over zero-shot inference across all six benchmark datasets and all pre-trained backbones. Zero-shot performance varies considerably across models and datasets, often remaining below 40% on multiple benchmarks. After introducing CSB, Macro-F1 scores increase markedly across all backbone–dataset combinations, with the majority of results exceeding 70%, and many reaching above 80%.

These gains are observed across both general-purpose models (CLIP ViT-B/16 and ViT-L/14) and pathology-specialized models (BiomedCLIP and PLIP), indicating that the improvements are not confined to a specific pretraining corpus. Notably, even models with relatively weak zero-shot performance, such as MI-Zero, exhibit substantial performance gains after incorporating CSB, with increases of more than 50 percentage points on several datasets.

Overall, these findings demonstrate that CSB consistently enhances downstream pathological classification performance under data-limited supervision, and that the proposed framework adapts diverse pre-trained vision–language backbones in a stable and effective manner.

### 5.2. Adaptation-based Methods

We compare HistoSB-Net with representative lightweight adaptation approaches, including prompt-based methods (CoOp, CoCoOp), multi-modal prompt learning (MaPLe), and feature-level adapters (CLIP-Adapter, Tip-Adapter). Unless otherwise specified, we follow the official training protocols of each baseline.

For methods whose official implementations use shorter schedules (CoCoOp, MaPLe, and Tip-Adapter-F), we additionally report results re-trained for 50 epochs under our unified training setting to ensure fair comparison. The results are shown in Table 5.

**Table 5.**
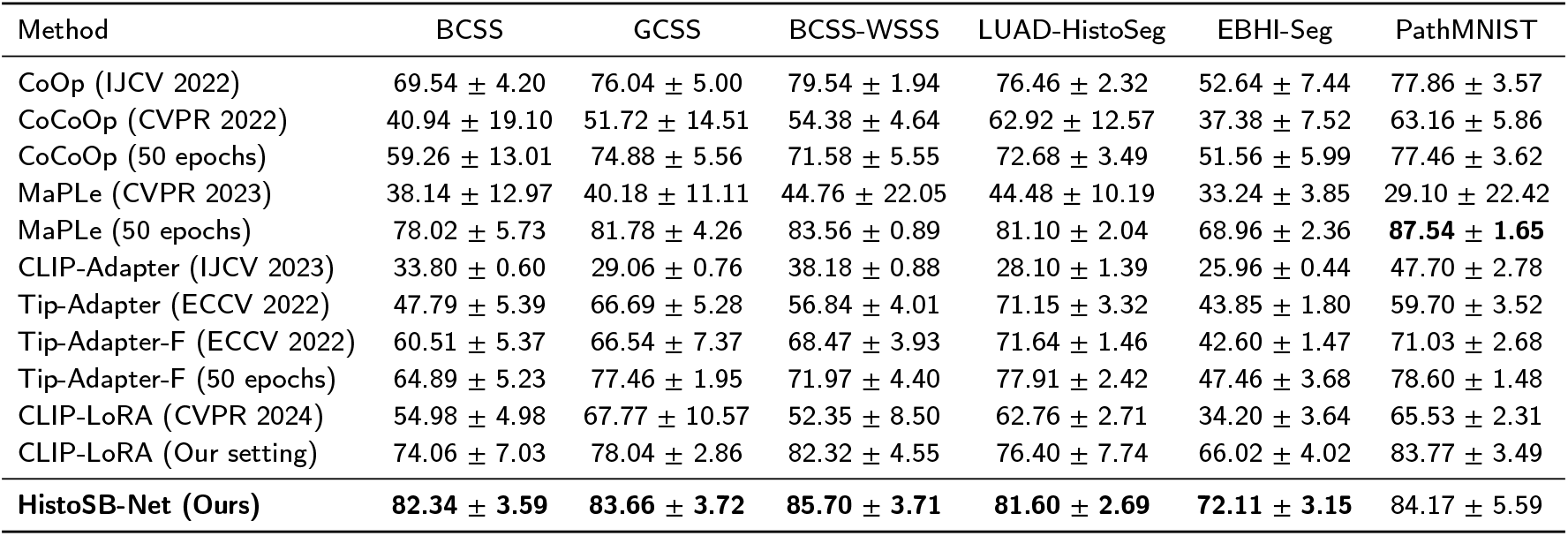
Macro-F1 (%) comparison of adaptation-based methods across pathology benchmarks.

As shown in Table 5, HistoSB-Net achieves the highest Macro-F1 scores on five out of six benchmarks (BCSS, GCSS, BCSS-WSSS, LUAD-HistoSeg, and EBHI-Seg), and remains competitive on PathMNIST, where MaPLe (50 epochs) attains the best result. This demonstrates that the proposed framework performs favorably against representative prompt-based and adapter-based adaptation methods under the same data-limited setting.

Prompt learning approaches such as CoOp and CoCoOp exhibit noticeable performance variability across datasets. In particular, CoCoOp shows relatively large standard deviations under the official training schedule, indicating sensitivity to stochastic initialization. Although extended 50-epoch training improves its stability and average performance, the results remain dataset-dependent. Similarly, MaPLe benefits substantially from longer training, achieving strong results on several benchmarks, yet its performance fluctuates more markedly across datasets compared with HistoSB-Net.

Feature-level adapters, including CLIP-Adapter and Tip-Adapter variants, generally provide moderate improvements but do not consistently reach the top performance tier. Even with extended training, Tip-Adapter-F remains below HistoSB-Net on most datasets.

For CLIP-LoRA, we report both the original training configuration from the source paper and results re-trained under our unified experimental setting. Under the official configuration, CLIP-LoRA exhibits a noticeable performance gap compared with HistoSB-Net across pathology benchmarks. Considering the parameter sensitivity induced by domain shift, we further re-train CLIP-LoRA using the same optimizer, learning rate schedule, dropout configuration, and adaptation hyperparameters as adopted for HistoSB-Net, leading to substantial performance improvements on all datasets.

This observation corroborates our earlier discussion that projection-level weight reparameterization is highly sensitive to optimization settings under domain-shifted pathology tasks. In contrast, all results of HistoSB-Net are obtained under a single unified training configuration, consistently outperforming CLIP-LoRA on five out of six benchmarks and achieving stronger cross-dataset consistency. These findings suggest that explicitly conditioning residual modulation on the frozen projection geometry provides a more stable adaptation mechanism than low-rank weight reparameterization alone.

### 5.3. Ablation Study

We analyze the scaling behavior and structural design of CSB through controlled ablations on supervision scale, encoder branch selection, and projection insertion strategy. Backbone architectures and evaluation metrics remain fixed unless otherwise specified.

#### 5.3.1. Effect of Supervision Scale

To evaluate behavior under varying supervision regimes, we vary the number of labeled samples per class (2, 4, 8, 16, and 32) and report corresponding Macro-F1 scores. This experiment examines whether CSB provides consistent gains across different levels of supervision.

As shown in Fig. 4, HistoSB-Net demonstrates a clear and monotonic performance improvement as the number of labeled samples per class increases from 2 to 32. The average Macro-F1 rises from 54.94% (2-shot) to 65.46% (4-shot), 74.23% (8-shot), 81.60% (16-shot), and 86.24% (32-shot). This consistent upward trend is observed across all six benchmarks, indicating stable scaling behavior under progressively richer supervision.

**Figure 4.**
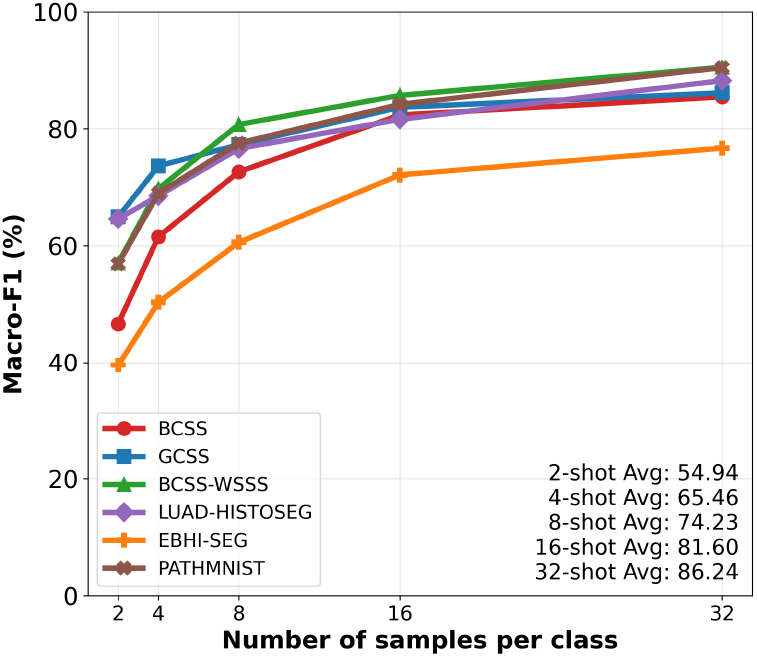
Performance scaling of HistoSB-Net with increasing supervision across six pathology benchmarks.

Importantly, no performance degradation is observed as supervision increases, indicating stable optimization behavior across low- and moderate-shot settings. These results confirm that the proposed framework maintains effectiveness across a wide range of supervision regimes.

#### 5.3.2. Encoder Branch Selection

This ablation examines how regulating different encoder branches affects image–text alignment and diagnostic performance. Specifically, we evaluate three configurations: applying CSB to the vision encoder only, the text encoder only, or both branches simultaneously. By isolating branch-specific modulation, this analysis reveals how semantic regulation in each modality contributes to cross-modal consistency and classification robustness.

Figure 5 presents the impact of applying CSB to different encoder branches. On average across datasets, applying CSB to the vision encoder achieves 81.53% Macro-F1, while applying it to the text encoder alone yields 75.21%. Joint modulation of both encoders attains the highest average performance at 81.60%.

**Figure 5.**
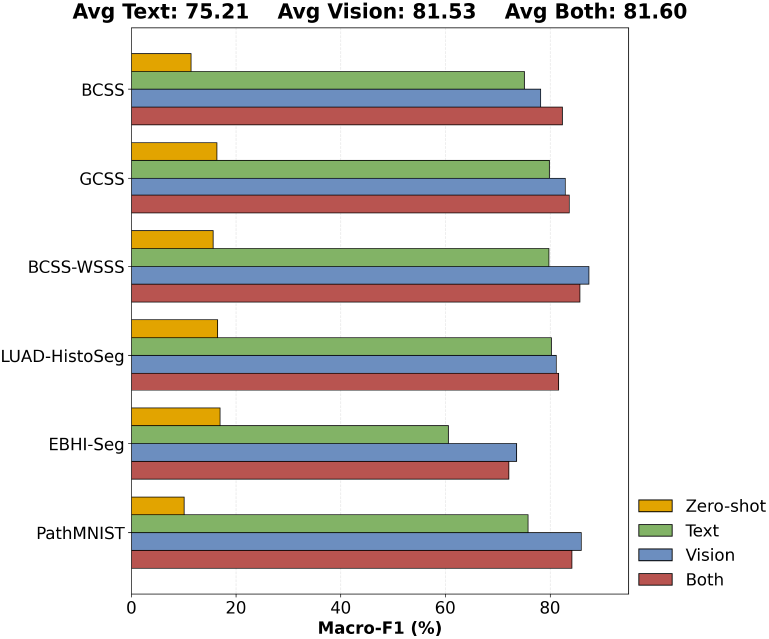
Effect of branch-specific CSB modulation on diagnostic performance across vision-only, text-only, and joint encoder configurations.

These results indicate that regulating the vision encoder contributes more substantially to downstream classification performance than text-only modulation. However, the slight improvement observed when jointly regulating both encoders suggests that simultaneous modulation of the vision and text encoders provides complementary cross-modal signals that extend beyond single-branch regulation. Furthermore, all branch-specific configurations substantially outperform the CLIP ViT-B/16 zero-shot baseline across all six benchmarks, demonstrating that projection-level regulation provides significant gains beyond frozen pre-trained representations.

#### 5.3.3. Projection Insertion Strategy

The influence of inserting CSB into different self-attention projections is examined to understand its impact on semantic alignment and diagnostic performance. CSB is applied separately to the Q, K, V, and O projections, as well as jointly to QKV, enabling a systematic comparison of distinct insertion strategies. The relative performance change is computed as the difference in Macro-F1 between each projection configuration and the query-only (Q) baseline, averaged across all evaluated datasets.

As illustrated in Fig. 6, different projection insertion strategies lead to distinct performance variations relative to the Q-only baseline. Inserting CSB into the K projection results in a performance decrease of -1.37%, whereas modulating the O and V projections yields moderate improvements of +0.55% and +0.73%, respectively. Jointly inserting CSB into all three projections (QKV) achieves the largest gain of +1.15%, representing the strongest empirical performance among the tested strategies.

**Figure 6.**
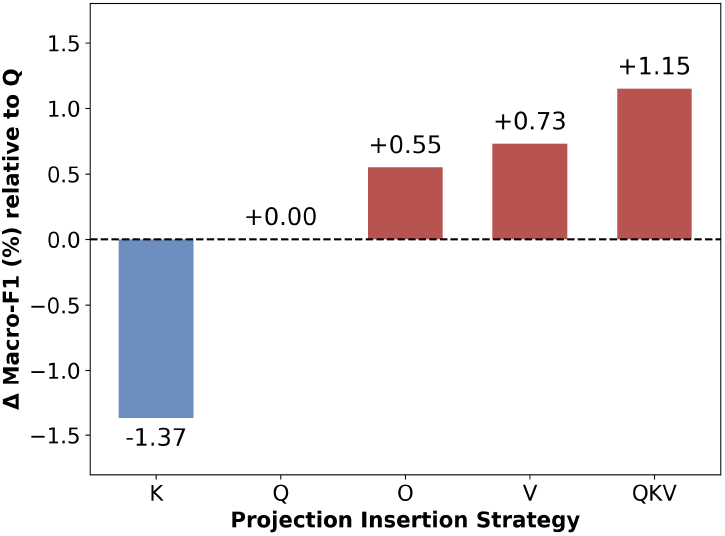
Relative Macro-F1 (%) change of CSB projection insertion strategies with respect to the query-only baseline, averaged across all datasets. Positive values denote gains and negative values denote drops.

These findings suggest that different attention projections contribute unequally to semantic regulation, and that coordinated modulation across multiple projections offers a more reliable configuration for enhancing downstream classification performance.

### 5.4. Analysis of Class Discriminability

A central question raised in the introduction is whether HistoSB-Net effectively alleviates intra-class heterogeneity and inter-class homogeneity in histopathological representation learning. To answer this question, we analyze class discriminability at both the representation and prediction levels through a prototype-based structural analysis and a class-level confusion analysis.

#### 5.4.1. Prototype-Based Margin Analysis

Let **x**_*i*_ ∈ ℝ^*d*^ denote the normalized embedding of sample *i*, and let *y*_*i*_ ∈ {1, …, *C*} be its ground-truth class label. For each class *k*, we define a class prototype as the mean embedding

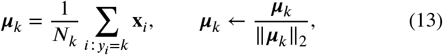

where *N*_*k*_ is the number of samples in class *k*. Similarity is measured by cosine similarity,

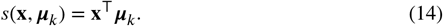

For each sample **x**_*i*_, the class discriminability margin is defined as

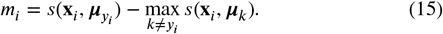

A positive margin indicates that the sample is closer to its own class prototype than to any competing class. We further compute the proportion of samples with positive margins,

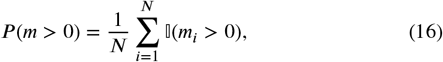

which reflects the fraction of correctly separated representations in the embedding space.

As shown in Fig. 7, HistoSB-Net consistently shifts the margin distributions toward larger positive values across all benchmark datasets. For BCSS, the mean margin increases from 0.010 (zero-shot) to 0.083, and the proportion of positive margins rises from 79.8% to 88.2%. Similar improvements are observed for GCSS (0.007 → 0.098; 81.9% → 90.4%) and BCSS-WSSS (0.010 → 0.080; 81.7% → 91.5%).

**Figure 7.**
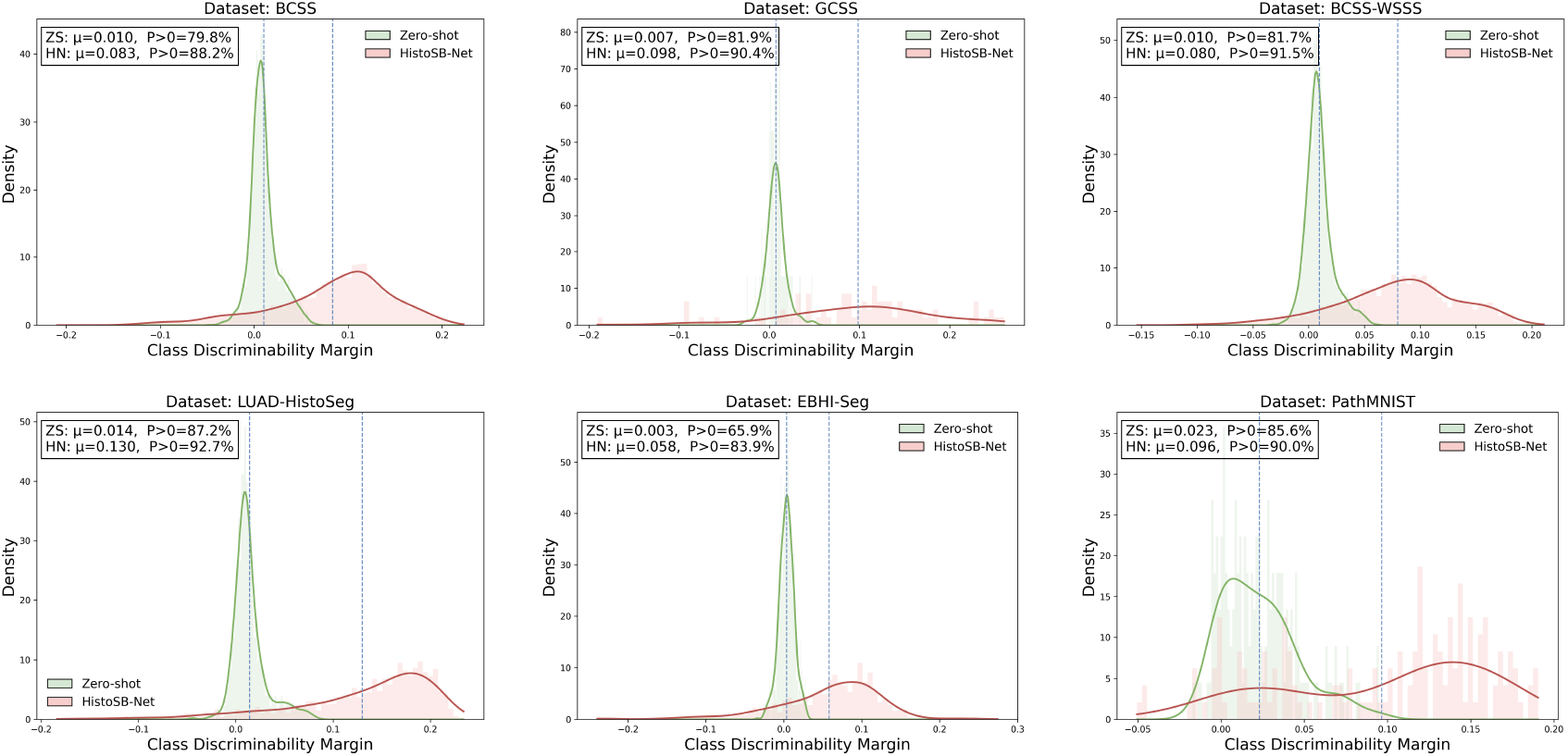
Class discriminability margin distributions across all benchmark datasets.

More pronounced expansion of the margin distribution is observed in LUAD-HistoSeg (0.014 → 0.130; 87.2% → 92.7%) and EBHI-Seg (0.003 → 0.058; 65.9% → 83.9%), indicating substantial separability gains on datasets that previously exhibited smaller margins. PathMNIST also shows consistent improvement, with the mean margin increasing from 0.023 to 0.096 and *P* (*m* > 0) rising from 85.6% to 90.0%.

Across all datasets, both the average margin and the proportion of positively separated samples increase in HistoSB-Net, demonstrating enhanced intra-class compactness and enlarged inter-class separation in the learned embedding space.

#### 5.4.2. Confusion Matrix Analysis

While margin distributions characterize geometric separability in the embedding space, it is equally important to examine how these structural improvements translate into class-level prediction behavior. To this end, we analyze the confusion matrices of zero-shot CLIP and HistoSB-Net across all datasets.

As shown in Fig. 8, zero-shot models exhibit noticeable off-diagonal activations across multiple datasets, indicating substantial inter-class overlap in the frozen representation space. After incorporating CSB, the adapted models consistently display stronger diagonal dominance and visibly reduced off-diagonal responses.

**Figure 8.**
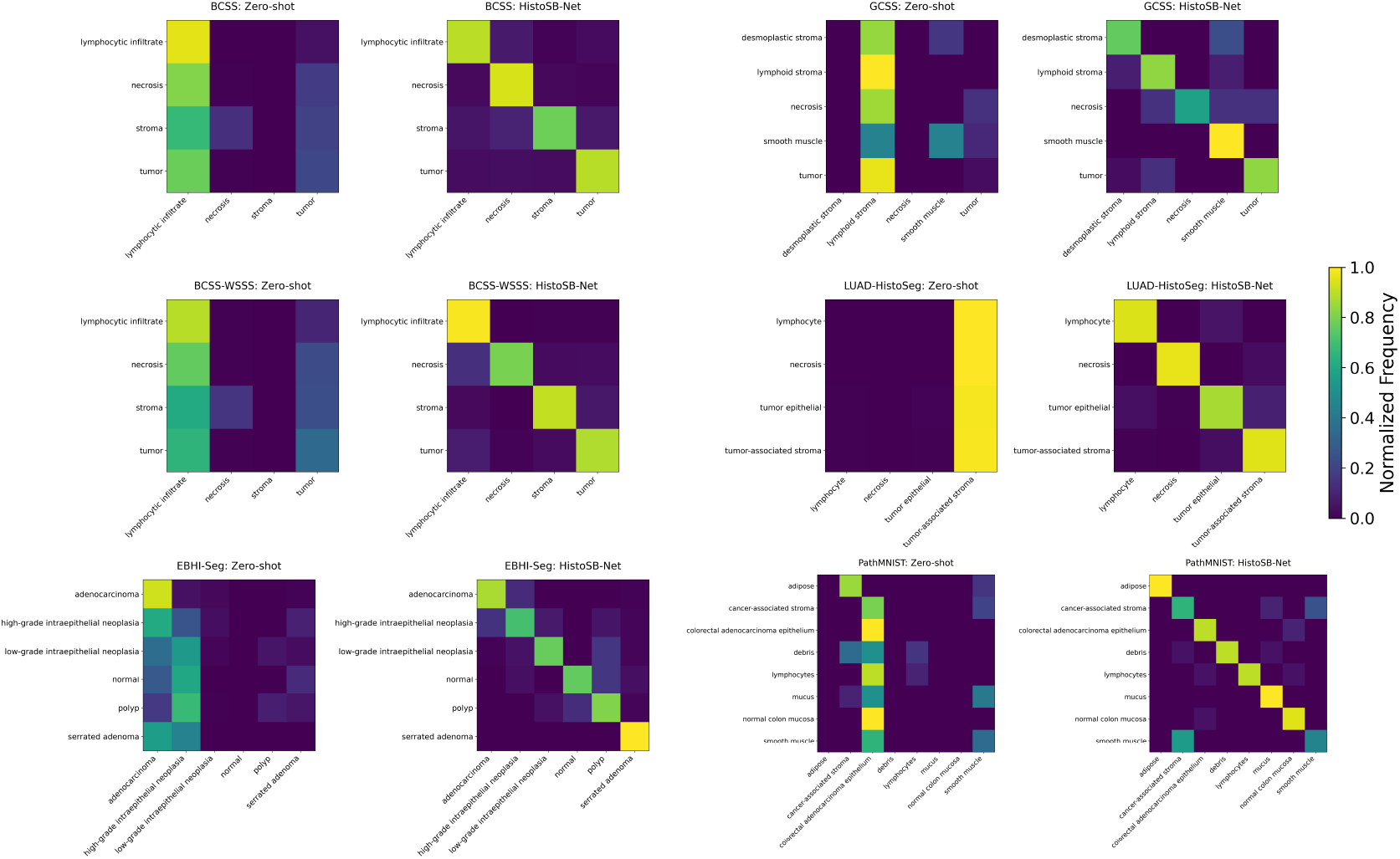
Confusion matrices across all benchmark datasets, comparing zero-shot CLIP ViT-B/16 with HistoSB-Net.

For example, in BCSS and BCSS-WSSS, stromal and tumor-related categories that previously showed substantial overlap become more distinctly separated. In EBHI-Seg and PathMNIST, the zero-shot baseline exhibits concentrated misclassification patterns, whereas HistoSB-Net produces more balanced and diagonally concentrated predictions.

These observations are consistent with the margin analysis results and suggest that the performance gains of HistoSB-Net are associated with improved representation geometry rather than solely decision-level recalibration.

### 5.5. Computational Cost and Parameter Efficiency

To evaluate the computational overhead of the proposed method, we report the total training time over 50 epochs, metpeak GPU memory consumption, and GPU utilization.

As summarized in Table 6, HistoSB-Net exhibits stable computational cost across benchmarks, with total training time over 50 epochs ranging from 37.40 s (GCSS) to 48.00 s (PathMNIST). Peak GPU memory consumption varies between 3394 MB and 5500 MB, corresponding to 13.82%– 22.39% of the memory capacity of an NVIDIA RTX 4090 GPU. The number of trainable parameters accounts for only 0.49% of the full ViT-B/16 backbone, corresponding to 0.74M trainable parameters out of 150.4M total parameters, confirming the parameter-efficient nature of the proposed CSB module. These results indicate that CSB introduces limited computational overhead while maintaining practical training efficiency under standard hardware conditions.

**Table 6.** Computational cost of training. Time (s) denotes the total training time over 50 epochs, and GPU Usage (%) is calculated based on the 24 GB memory capacity of an NVIDIA RTX 4090 GPU.

| Dataset | Time (s) | GPU (MB) | Usage (%) |
| --- | --- | --- | --- |
| PathMNIST | 48.00 | 4440.25 | 18.08 |
| BCSS | 45.80 | 5483.05 | 22.32 |
| BCSS-WSSS | 42.20 | 5482.25 | 22.32 |
| GCSS | 37.40 | 3394.25 | 13.82 |
| LUAD-HistoSeg | 38.20 | 5478.25 | 22.30 |
| EBHI-Seg | 42.80 | 5500.25 | 22.39 |

## 6. Conclusion

This work presented HistoSB-Net, a semantic bridging framework for adapting pre-trained VLMs to multimodal histopathological diagnosis under data-limited supervision. The study identified semantic misalignment between natural image–text pretraining and pathology-specific diagnostic concepts as a key barrier to effective transfer, and addressed this issue through CSB within the self-attention projection space of both vision and text encoders.

Rather than employing explicit cross-attention or full fine-tuning, the proposed CSB module modulates frozen attention projections through a lightweight nonlinear bottleneck, preserving the backbone architecture while introducing structured projection-level regulation. The resulting model maintains strong parameter efficiency, requiring only 0.49% of trainable parameters relative to the full backbone and incurring moderate computational overhead.

Extensive experiments across six pathology benchmarks demonstrate consistent and substantial improvements over zero-shot inference and representative adaptation baselines under few-shot supervision. In addition to improved Macro-F1 performance across diverse pre-trained backbones, controlled ablation studies confirm the contributions of supervision scale, encoder branch selection, and projection insertion strategy. Representation-level analyses further show enlarged class discriminability margins and strengthened diagonal dominance in confusion matrices, indicating enhanced intra-class compactness and inter-class separation in the learned embedding space.

Several directions remain for future investigation. Extending projection-level semantic bridging to hierarchical or multi-scale VLM architectures may further enhance adaptability to complex histopathological structures. Incorporating domain-aware textual priors could introduce additional semantic regularization within the shared embedding space. Moreover, integrating automated global WSI handling directly into the learning pipeline may provide a more unified and scalable solution for large-scale digital pathology.

Overall, these findings suggest that regulating attention projections constitutes an effective and computationally manageable pathway for transferring pre-trained VLMs to data-limited medical imaging domains.

